## Supplementary Information for "Protein corona formed on lipid nanoparticles compromises delivery efficiency of mRNA cargo"

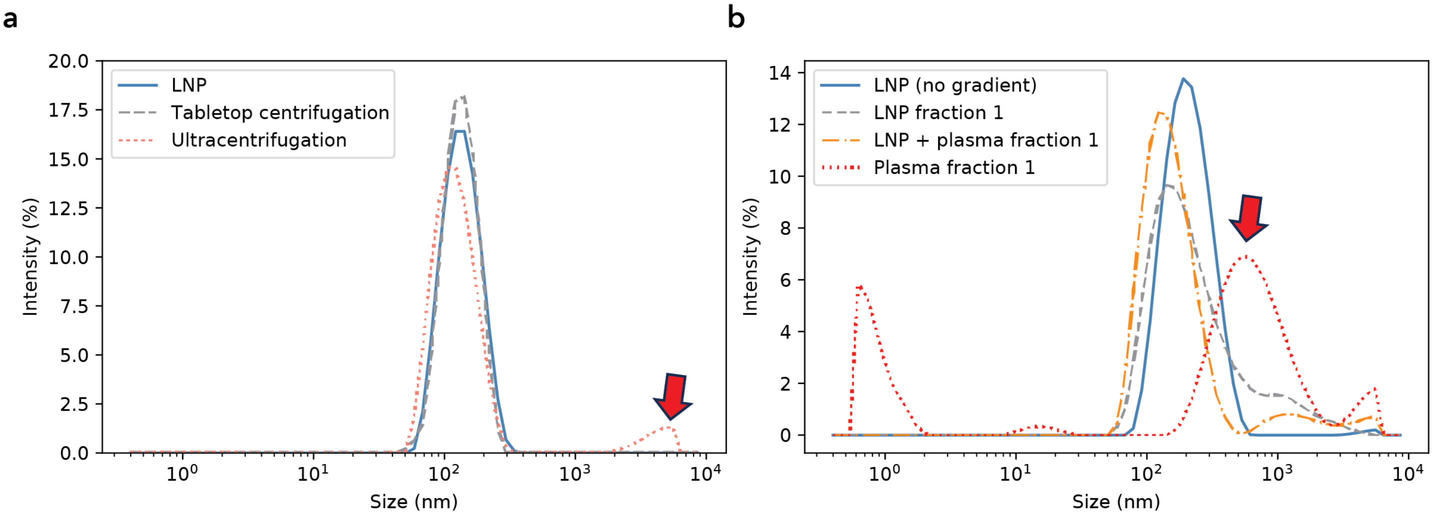


**Supplementary Figure 1. DLS shows limitations of current techniques to isolate the protein corona on lipid nanoparticles.** (a) DLS of LNPs in the supernatant before (blue line) and after tabletop centrifugation (dotted grey line) and ultracentrifugation (dotted orange line) reveal lack of pelleting and aggregation (highlighted by red arrow), respectively. (b) 1-mL fractions are collected top to bottom from prepared three-layer (30%, 15%, 0%) iodixanol density gradients of LNPs alone, LNPs incubated with plasma, and plasma alone centrifuged for 3 hours. DLS of current strategies for gradient layering for ultracentrifugation show that the native biological particles in the plasma control gradient (red arrow) are present in the parallel first fraction for the LNP gradient.

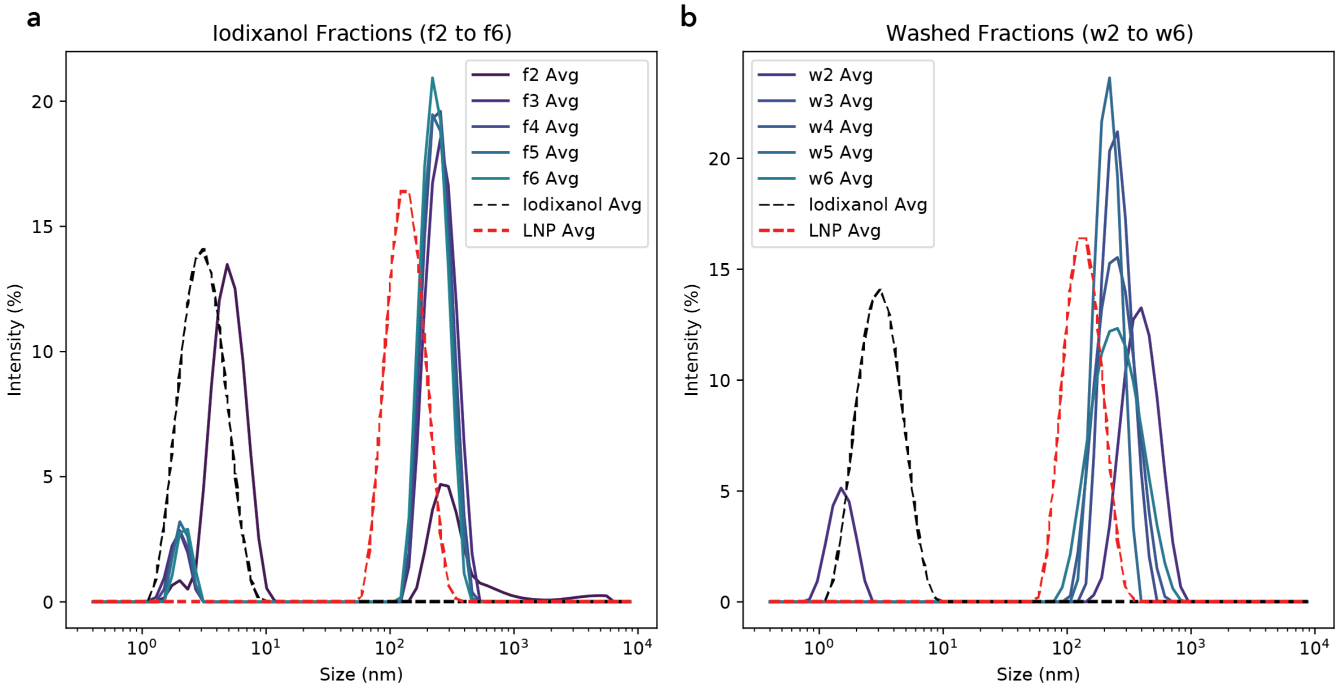


**Supplementary Figure 2. DLS of particles after density gradient centrifugation (DGC)**. The LNPs were loaded into a density gradient and fractions were collected top to bottom according to the method used for proteomics characterization. To assess the stability of particles after centrifugation, LNP size was characterized via DLS in (a) the iodixanol medium and (b) after washing. The fractions show similar sized LNPs to the control LNPs that did not undergo DGC (in red) with an additional peak present where iodixanol forms small particles in solution (dotted black line). The shift in size to the right is likely due to differences in solution properties. These measurements can only qualitatively confirm particle preservation because the refractive index and viscosity of the iodixanol changes within the gradient, affecting the DLS size calculations. The DLS calculation relies on the assumption that the particles are in a solution with the same refractive index and viscosity for comparison. To address this problem, we washed samples selected for proteomics characterization twice in PBS in 3-kDa MWCO Amicon centrifugal filters to remove the iodixanol medium. The washed particles also show similar size profiles to the control LNPs in red.

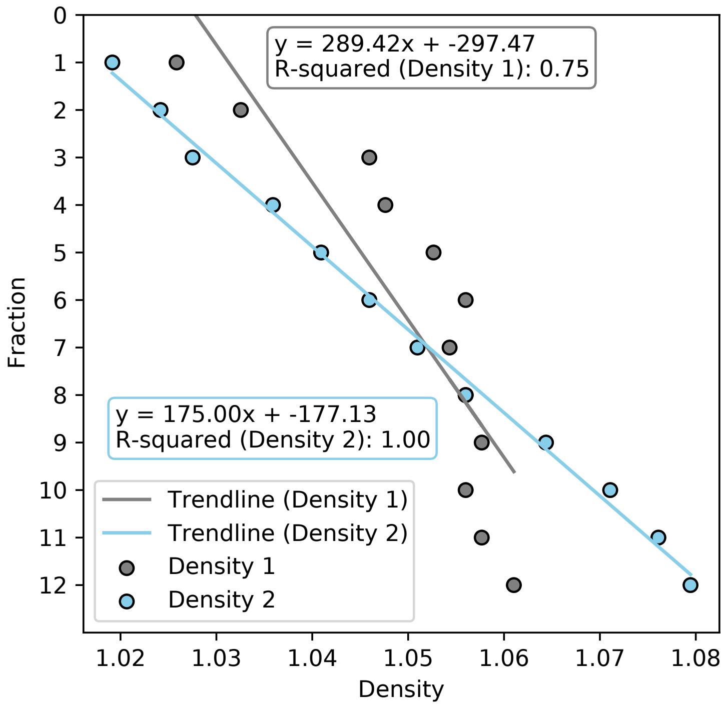


**Supplementary Figure 3. Density linearity within the gradient.** The density is calculated by measuring refractive index via refractometer and converting based on known standard curves for iodixanol. The density change within the first 12 fractions of the gradient must be linear to ensure appropriate separations. As shown above, bumping during gradient loading may result in a poor linear trend (gray line; Density 1, $R^2 = 0.75$). In contrast, careful layering will result in a linear density gradient (blue line; Density 2, $R^2 = 1.00$). These measurements highlight the need for quality control during gradient loading. Loading these gradients is technically challenging and overlooking quality control in density distribution throughout the tube may lead to LNP sample contamination and therefore misleading protein corona results.

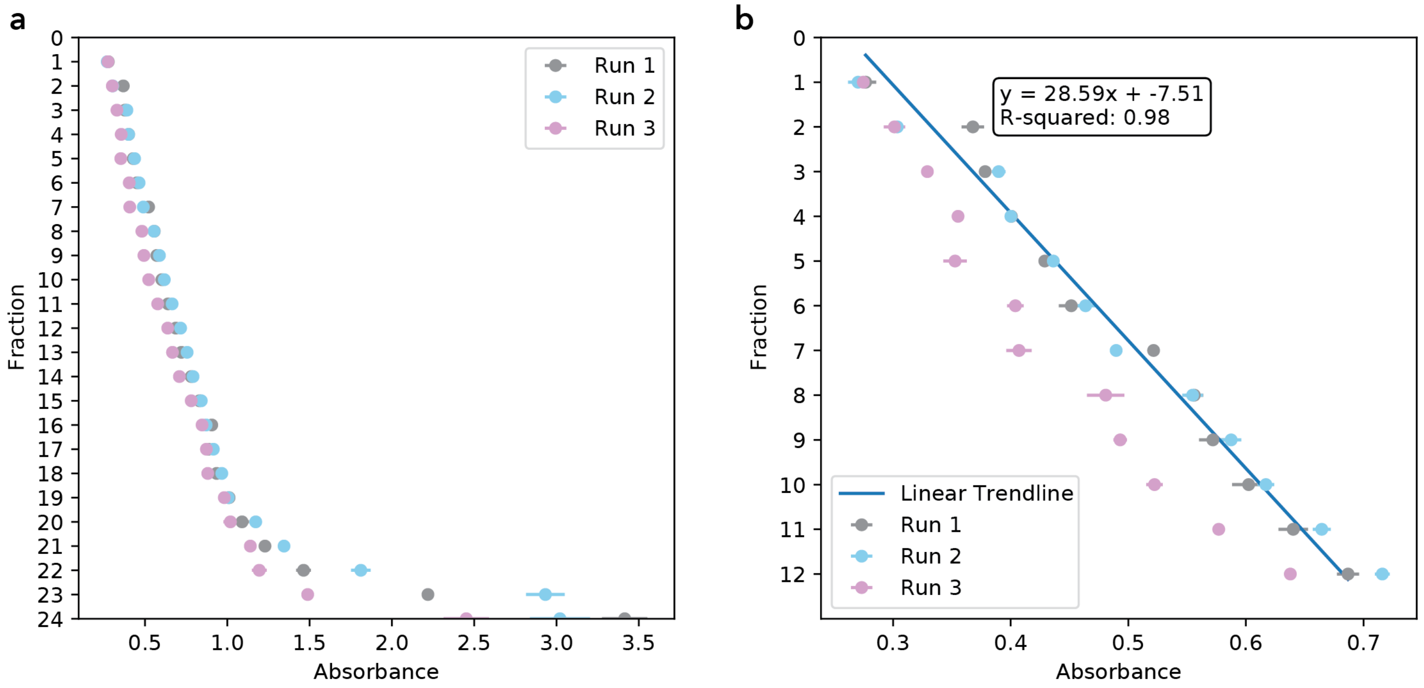


**Supplementary Figure 4. Absorbance linearity within the gradient.** (a) The absorbance at wavelength 340 nm, the peak absorbance for OptiPrep, was measured for fractions after DGC. (b) As shown by the inset, the absorbance shows a linear trend within the fractions of interest. These absorbance measurements are a quick method for confirming successful gradient preparation, such as Run 2 with a linear trendline ($R^2 = 0.99$).

**Supplementary Table 1. LNP localization within the density gradient.**

| Experiment # | Area under the curve (%) |
|---|---|
| Run 1 | 48.62 |
| Run 2 | 72.25 |
| Run 3 | 81.70 |

We estimated the percentage of LNPs within the peak by calculating the area under the measured fluorescence curve (Fig. 2c) for each run between fractions 2-6 relative to the total area under the measured fluorescence curve using the trapezoidal rule (trapz function from scipy.integrate). A baseline area, connected by the first and last data points, is subtracted to adjust the baseline to zero. The average was determined to be 67.52 with a standard deviation of 13.91.

**Supplementary Table 2. Relative abundance of serum albumin in upper fractions of density gradient.**

| | **Serum albumin relative abundance (%)** | |
|---|---|---|
| *Method* | *LNP sample* | *Plasma control sample* |
| **Method 1**<br>4 hour, 40 k rpm, 4 °C, layers of 15% and 30% iodixanol | 90.88 | 70.26 |
| **Method 2**<br>16 hour, 36 k rpm, 4 °C, layers of 30%, 25%, 20%, 15%, 10%, and 5% iodixanol | 24.12 | 21.22 |

Supplementary Table 2 shows the decrease in relative corona abundance for the most blood plasma-abundant protein, serum albumin. This suggests an overall improvement in the separation of free proteins from the fractions containing the LNPs. The relative abundance for the method 1 was calculated by determining the average relative abundance across fractions 2-6, which is the equivalent of pooled fractions 2-6 in method 2.

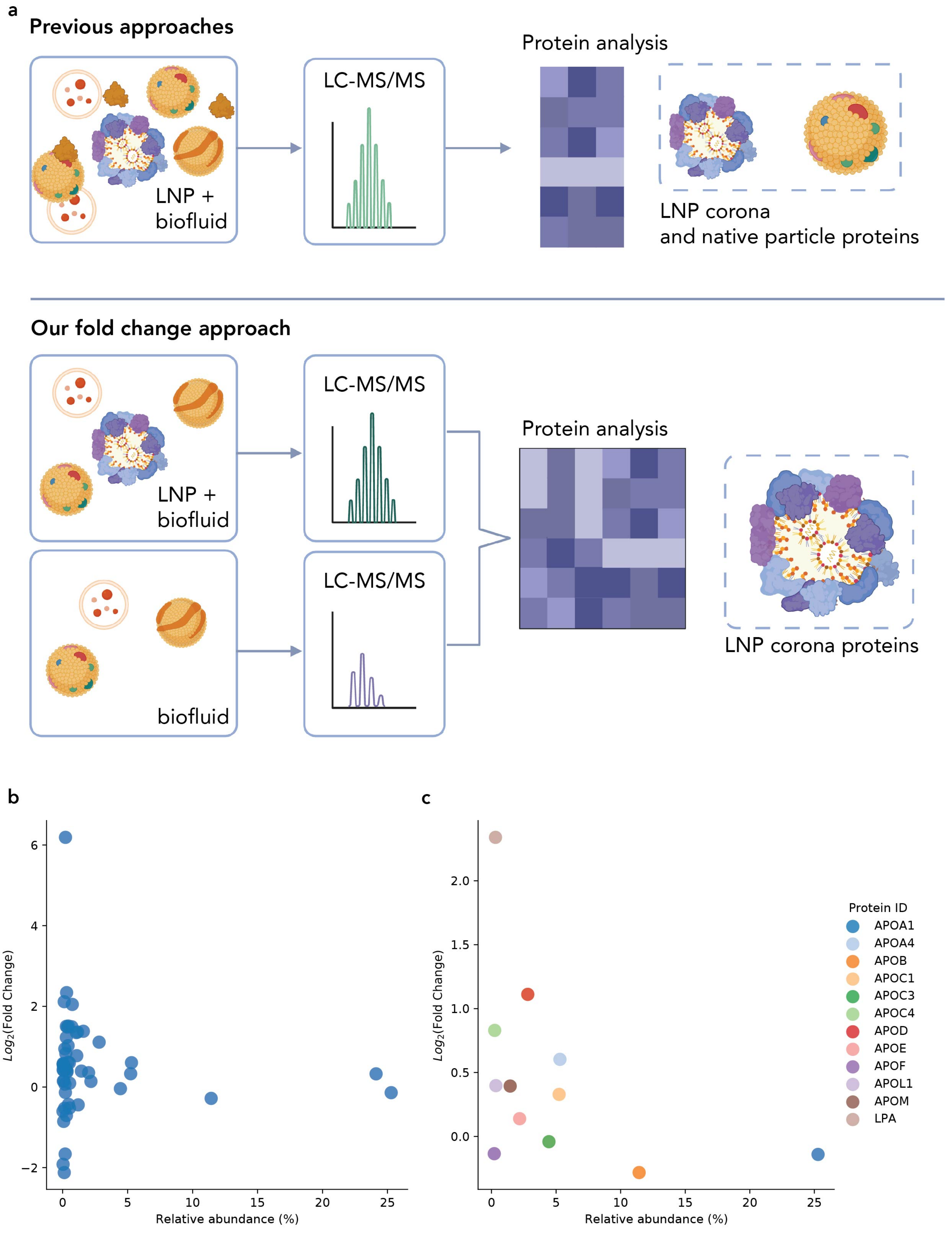
a
Previous approaches
LNP +
biofluid
LC-MS/MS
Protein analysis
LNP corona
and native particle proteins
Our fold change approach
LNP +
biofluid
biofluid
LC-MS/MS
LC-MS/MS
Protein analysis
LNP corona proteins
b
$Log_2$(Fold Change)
Relative abundance (%)
c
$Log_2$(Fold Change)
Relative abundance (%)
Protein ID
APOA1
APOA4
APOB
APOC1
APOC3
APOC4
APOD
APOE
APOF
APOL1
APOM
LPA

**Supplementary Figure 5. Comparison of logarithmic fold-change against relative abundance (%) for identified lipoproteins.** (a) A comparison of previous approaches which examine relative abundance of only the LNP sample and our approach which quantifies differences between the LNP sample and a biofluid control. The scatter plots show the correlation between the data analysis workflow (fold change relative to plasma) suggested in this paper and previous approaches (relative abundance (%)) for (b) all identified proteins and (c) apolipoproteins. The Pearson correlation coefficient for all proteins and lipoproteins was calculated to be -0.0871 and -0.426, respectively. The negative correlation between the two approaches highlights the need for more controlled approaches to analyzing the lipoprotein-LNP interactions.

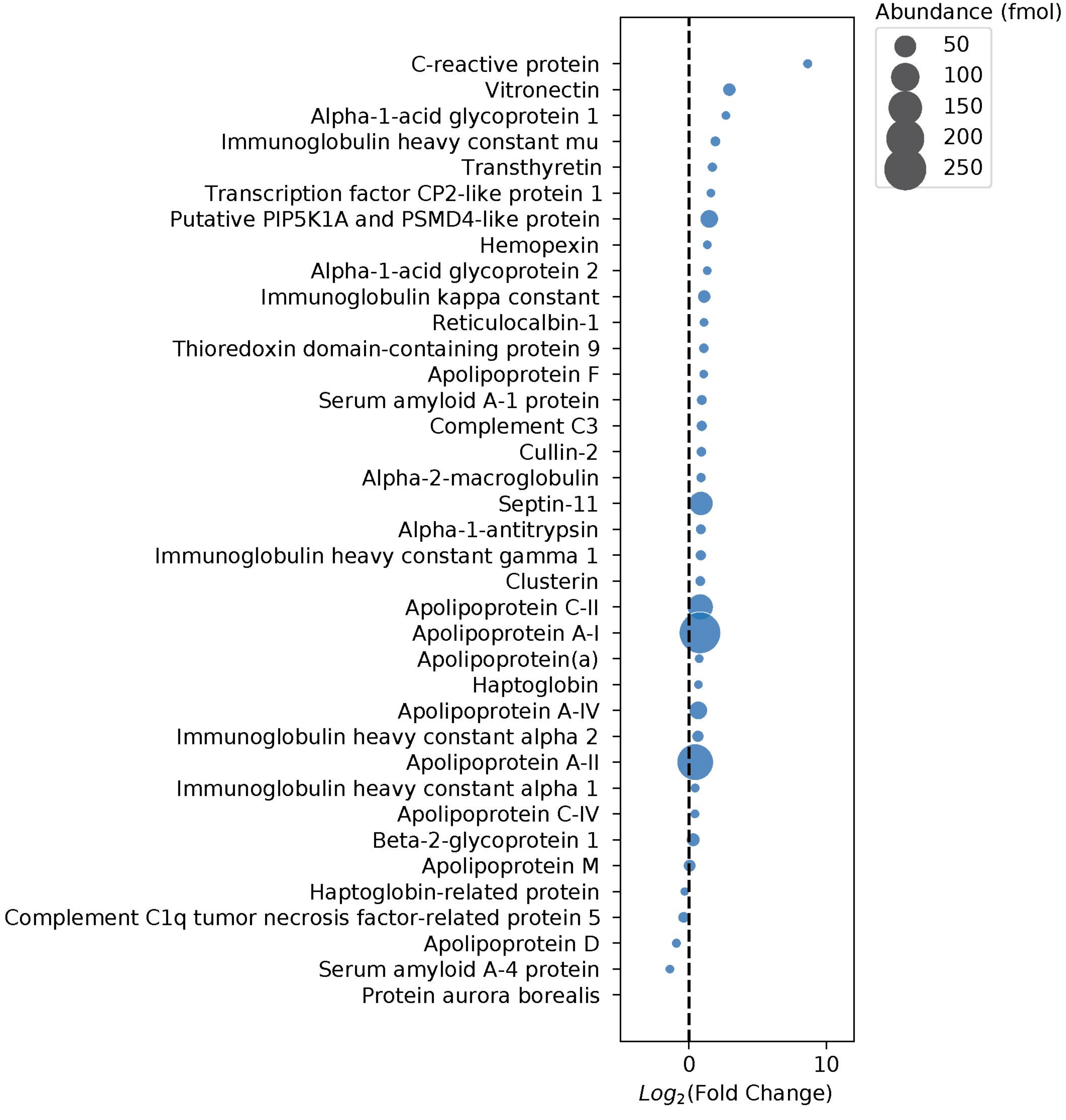


**Supplementary Figure 6. LC-MS/MS of samples processed in parallel.** Logarithmic fold change of proteins, with bubble size showing abundance (fmol) of samples processed in parallel. The protein abundance for protein-LNP samples was analyzed relative to plasma control fractions and was filtered for adjusted p-value (q-value) <0.05. C-reactive protein and vitronectin were found to have high enrichment, in agreement with other datasets we collected. However, ApoE is not enriched across the parallel experiments, despite prior reports suggesting that ApoE adsorption drives downstream behavior including cell uptake.[1] Independent LC-MS/MS processing experiments show that differences in ApoE abundance (fmol) between the control sample and the LNP are not statistically different.

**Supplementary Table 3. Proteins enriched across three independent experiments**.

| Protein | $Log_2$(fold change) | STDV |
|---|---|---|
| Alpha-2-macroglobulin | 1.31 | 0.42 |
| Apolipoprotein(a) | 2.52 | 0.50 |
| C-reactive protein | 8.47 | 2.07 |
| Haptoglobin-related protein | 1.09 | 0.62 |
| Vitronectin | 2.33 | 0.25 |

**Supplementary Table 4. Estimate of protein mass relative to mRNA mass based on LC-MS/MS abundances.**

| Protein | Abundance detected via LC-MS/MS (fmol) | MW (g/mol) | Mass of protein (ng) | Ratio of mRNA to protein |
|---|---|---|---|---|
| **CRP** | 4.04 | 25039 | 0.10 | 1.98E+05 |
| **VTN** | 13.74 | 54306 | 0.75 | 2.68E+04 |
| **A2M** | 5.25 | 163291 | 0.86 | 2.33E+04 |
| **ApoE** | 40.25 | 36154 | 1.46 | 1.37E+04 |

**Supplementary Table 5. Estimate of protein concentrations for selected proteins in native human plasma.**

| Protein | Native human plasma concentration (mg/mL) |
|---|---|
| **CRP** | 0.009 (ref)[2] |
| **VTN** | 0.02 (ref)[3] |
| **A2M** | 178 (ref)[4] |
| **ApoE** | 0.05 (ref)[5] |

Note: The total concentration of protein in solution for protein incubations was 0.01 mg/mL (*2 µg in 200 µL of media*).

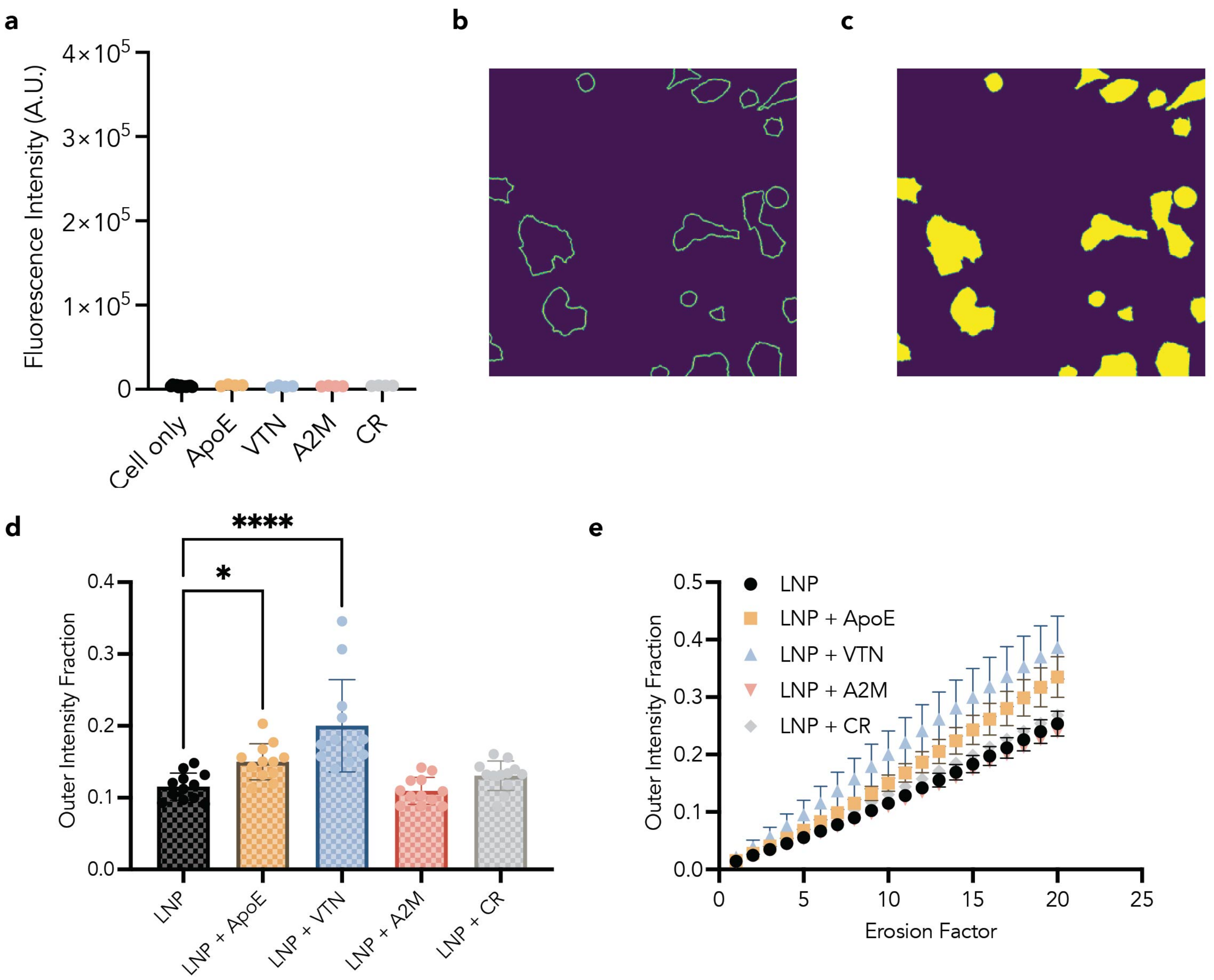


**Supplementary Figure 7. Cy5 signal cellular localization analysis.** (a) Cy5 fluorescence intensity per cell of cells incubated with protein show no significant signal. To compare the position of the LNPs associated with the cell, (b) the signal in the outer regions of the cells shown in green relative to (c) the rest of the cell membrane shown in yellow was analyzed. (d) Localization analysis reveals that the VTN-LNP complexes had more signal in the outer region of the cell in comparison to LNPs alone. To compare the position of the LNPs associated with the cell, the signal in the outer regions of the cells relative to the rest of the cell membrane was analyzed across different levels of erosion. (e) These changes in outer intensity hold over higher rates of erosion. N = 4 technical replicates, n = 3 biological replicates. Data points shown are 3 averaged FOV for each technical replicate. Error bars all denote standard deviation, One-way ANOVA test where * and **** represent $p \leq 0.05$ and $p \leq 0.0001$ respectively.

a b

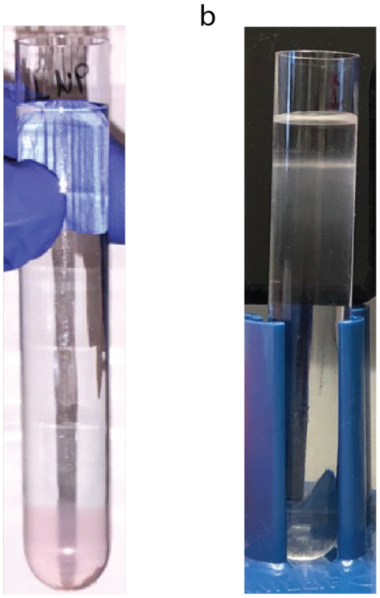


**Supplementary Figure 8. Linearity within the gradient.** (a) The gradient should be loaded such that the layers are visually distinguishable due to their differences in refractive index. The LNPs (here tagged with a dye for visualization) are loaded on the bottom of the gradient. (b) After centrifugation, the LNPs (undyed) should partition throughout the tube according to their density.
